# CDK5 Inhibitor Seliciclib Promotes Osteoblastic Differentiation of MSCs and Suppresses the Migration of MG-63 Osteosarcoma Cells

**DOI:** 10.1101/2020.12.07.415612

**Authors:** Hong Fu, Haoyu Zhao, Yali Yang, Ke Duan, Tailin Guo

## Abstract

CDK5 belongs to the cycling dependent kinase family, which is multifunctional and plays an important role in neural differentiation. However, the role of CDK5 in osteoblastic differentiation remains unclear. The present study investigated functions and molecular mechanism of CDK5 in osteoblastic differentiation. It was found that, the addition of CDK5 inhibitor Seliciclib promoted the expression of *Runx2*, *ALP*, *OCN* and *OPN* of MSCs and the mineralization of MC-3T3E1 cells. Seliciclib enhanced the development of F-actin, nuclear localization of β-catenin and YAP, as well as the expression of RMRP RNA. When F-actin was suppressed by Blebbistatin, the nuclear localization of YAP and β-catenin, and expression of *RMRP RNA* as well as *Runx2* and *ALP* were decreased. These indicate Seliciclib promotes osteoblastic differentiation mainly by F-actin. Moreover, Seliciclib also suppressed the migration of MG-63, suggesting a potential application for Seliciclib in bone defect repair and inhibition of the migratiion of osteosarcoma cells.

## Introduction

CDK5 is a member of the cyclin-dependent kinase (CDK) family and shares 60% sequence identity to CDK1. Unlike other CDKs, CDK5 is the only CDK that does not play a role in the cell cycle. It is expressed ubiquitously and mostly expressed in the brain [1].

Importantly, CDK5 is an upstream regulator of F-actin dynamics, affecting it differently under different situations. P35/CDK5 kinase can phosphorylate the serine/threonine kinasep21 (Rac1)-activated kinase1 (Pak1) on site T212; this leads to its inactivation in neurons and differentiating epithelial cells [2, 3]. Pak1 is a positive regulator for RAC family small GTPase 1, a kinase able to promote the polymerization of actin. CDK5 can be a negative regulator of ras homolog family member A (RhoA). CDK5 phosphorylates DLC1 (deleted in liver cancer 1) on several sites (e.g., S120) to suppress RhoA [4]. DLC1 is aGTPase-activating protein (GAP) that can down-regulate the activation of RhoA [5]. CDK5 upregulates actin polymerization by phosphorylating, and thus suppressing, non-receptor tyrosine kinase Src. Src activates the Rho GTPase activating protein p190 (p190 RhoGAP), a suppressor of RhoA [6]. However the F-actin can promote the osteoblastic differentiation of MSCs [7]. The effects of CDK5 on F-actin dynamics in mesenchymal stem cells (MSCs) is unclear.

CDK5 also binds to β-catenin to reduce N-cadherin-mediated and calcium-dependent cell-cell adhesion [8]. Moreover, CDK5/p35 complex binds to β-catenin, thereby blocking its function as a transcriptian factor [9]. While during osteoblastic differentiation of MSCs, Wnt signaling pathways play important roles. The Wnt/β-catenin signaling pathway belongs to the canonical pathway. Beta-catenin has many functions important for osteoblastic differentiation. It is both a transcription factor and a membrane skeleton junction protein. It is involved in the mechanotransduction-mediated osteoblastic differentiation [10]. What’s more Yes-associated Protein (YAP) is a transcriptional factor regulated by the Hippo pathway. YAP can promote osteogenesis and suppress the adipogenic differentiation of MSCs by interacting with β-catenin and maintaining its nuclear localization [11]. Importantly, the nuclear localization of YAP can be promoted by the well-formed F-actin [12].

Of many discovered functions of CDK5, the most well-known one is its role in the normal development of cerebral cortex via promoting neuronal migration [13]. CDK5 regulates cell migration by phosphorylating the talinhead (TH) domain [14]. Phosphorylation of the TH domain helps the assembly of adhesion complexes, maintaining a balance between the assembly and disassembly of these complexes as well as a balanced lamellipodia stabilization-destabilization. Balance of lamellipodia stabilization-destabilization is needed for cell migration. Furthermore, as a result of its roles in cell migration and cell-cell adhesion, CDK5 is involved in the metastasis of many cancer [15–17]. Osteosarcoma, occurring frequently in adolescents, is the most common and highly metastatic primary bone tumor [18, 19]. Most cancer deaths are caused by metastases rather than by primary tumor growth. Finding drag to inhibite the metastasis of osteosarcoma is essential to treat osteosarcoma.

Considering the association of CDK5 with F-actin, β-catenin and cell migration, we hypothesized that CDK5 may play an important roles in the osteoblastic differentiation of MSCs and the migration of osteosarcoma cells. Therefore, this study was conducted to understand the roles of CDK5 in these processes.

## Materials and methods

### Cells and culture

MSCs were extracted from the bone marrow of Sprague-Dawley rat 2 weeks old. MSCs were cultured (95% relative humidity, 37℃, 5% CO2)in a proliferation medium [α-MEM, 10% fetal bovine serum (FBS), 1% antibiotic/antimycotic (all Hyclone)] with a medium change every 2 day.

MC-3T3 cells (ATCC, Virginia, US) were cultured in an osteogenic induction medium (1×10-7 mol/l dexamethasone, 1×10-2 mol/l β-glycerol phosphate disodium, 5×10-2 mm/l vitamin C) with a medium change every 2 day. MG-63 cells (ATCC, Virginia, US were cultured in a high-glucose medium [DMEM, 10% FBS, 1% antibiotic/antimycotic (all Hyclone)] with a medium change every 2 day.

### Cell plate and inhibitor use

For RT-qPCR, MSCs were seeded at 5×10^4^, 3×10^4^, 2×10^4^and 2×10^4^ cells/well in 12-plates and incubated for 1, 3, 5 and 7 d. For Alizarin red staining, MC-3T3E1 cells were seeded in a 24-well plate at 3×10^4^ cells/well and incubated for 14 days. For F-actin staining and immunofluorescent staining of YAP and β-Catenin, MSCs were seeded in a 48-well plate at 3×10^3^ cells/well and incubated for 5 days. For migration alalysis, MG-63 cells were seeded in a 24-well transwelldish (8μm bore diameter, Corning)at 2×10^4^ cells/well and incubated for 48h. Unless otherwise specified, the concentration of Seliciclib (CDK5 inhibitor, MedChemExpress)treated was 9 μM. The concentration of Blebbistatin (Ble, myosin II inhibitor, MedChemExpress)treated was 5 μM.

### RNA extraction and RT-qPCR

Cell were lyzed with TRIzol (100μl/well) (Invitrogen, Carlsbad, CA, USA) and transferred into an Eppendorf tube. After addition of 20 μl of chloroform, the tube was centrifuged (4℃, 1.2×10^4^ g, 15 min. The supernatant was collected, mixed with 50 μl of 2-isopropanol, allowed to rest for 10 min, and centrifuged (4℃, 1.2×10^4^ g, 10 min).The supernatant was discarded; pellet was mixed with 100 μl of 75% ethanol, centrifuged (4℃, 7.5×10^3^ g, 5 min). The pellet was dissolved in 20 μl of DNase/RNase-free water (Biosharp, GuangZhou, China). Then, the RNA was reverse transcribed to cDNA (Revertid First Strand cDNA Synthesis Kit, Bio-Rad, Hercules, CA, USA) and then amplified by real-time quantitative polymerase-chain reaction (RT-qPCR, SybrGreen PCR MasterMix Kit, Bio-Rad) using primers listed in Table 1. Relative gene expression was normalized to GAPDH expression. Throughout this study, each quantitative experiment included three parallel samples.

**Table 1.**
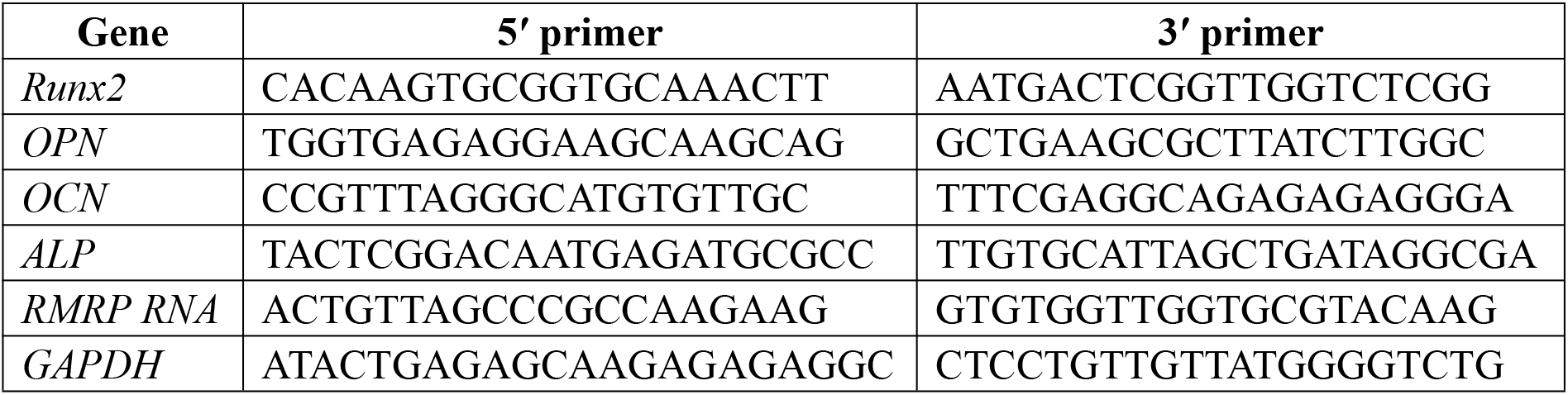
The sequence of RT-qPCR primer

### Immunofluorescent staining and F-actin staining

Cells were fixed for 3 h in PBS containing5% glutaraldehyde, permeabilized for 5 min in PBS supplemented with 0.1% Triton X-100 (Bioforxx,German), and blocked for 40 min in PBS containing 5% bovine serum albumin (BSA). They were probed with rabbit IgG to rat YAP (1:100, Cell Signaling Technology, Boston, USA)and β-catenin (1:300, Abcam, Cambridge Science Park, UK)(4℃, overnight in)and probed again with fluorescent-tagged goat IgG to rabbit IgG (1:500, Cell Signaling Technology; room temperature, 1 h), both in PBS supplemented with 1% BSA and 0.3% Triton X-100. Finally, nuclei were stained with DAPI (2 μg/ml, room temperature, 5 min, Sigma, St. Louis, Missouri, USA).

Additionally, after permeation with Triton X-100, some samples were stained for F-actin by treatment with 20 μg/ml rhodamine-conjugated phalloidin (room temperature, 40 min;Sigma, St. Louis, Missouri, USA) and stained with DAPI. All samples were observed under a fluorescence microscope (Vert.A1, Car Zeiss AG, Jena, Germany)and a confocal microscope (*/A1R+, Nikon, Tokyo, Japan).

### Alizarin red staining

MC-3T3 cells were fixed in 95% ethanol (20 min), incubated in 0.1% Alizarin red (Solarbio, for 20 minutes, and then observed under a stereo microscope (SMZ7 45, Nikon, Tokyo, Japan).

### Cell migration analysis

High-glucose DMEM (600 μl)supplemented with 10% FBS was added to the lower chamber of Transwell (Corning);100 μl of serum-free DMEM containing 1×10^5^ MG-63 cells was added to upper chamber. Subsequently, 10 μl of 1% BSA was added to the upper chamber to maintain an equivalent osmotic pressure. After incubation for 48 h, the upper chamber was treated with 4% paraformaldehyde for 20 min, stained with 0.1% crystal violet solution for 20 min, and observed under a microscope (Vert.A1, Car Zeiss AG, Jena, Germany). Five fields of view were randomly captured for cell counting.

### Statistical analysis

Data from three independent experiments were analyzed by independent t-test (GraphPad Prism 7, GraphPad, San Diego, CA, USA). A p-value <0.05 was considered statistically significant.

## Results

### Effects of Seliciclib on osteoblastic differentiation

On days 1, 5, and 7, the expression of *Runx2* was significantly higher in the Seliciclib-treated group than in the control group . On day 3, the difference between the two groups was negligible. The expression of *ALP* was significantly higher in the Seliciclib-treated group (vs. control) on days 1, 3 and 7. On day 5, the difference was minimal. The expression of *OCN* was significantly higher in the Seliciclib-treated group on days 1, 5, and 7, and was significantly lower in that group (both vs. control) on day 3. The expression of *OPN* was significantly higher in the Seliciclib-treated group than in the control on days 1 and 7, but was significantly lower on days 3 and 5. Overall, the relative expression of the four genes (i.e., Seliciclib-treated/control) generally increased between days 1–7 (Fig. 1A).*In vitro* mineralization is an indicator of osteoblastic differentiation. Alizarin red staining found that, after culture in the osteogenic induction medium for 14 d, the Seliciclib-treated group stained more intensely than did the control group (Fig. 1B).

**Figure 1.**
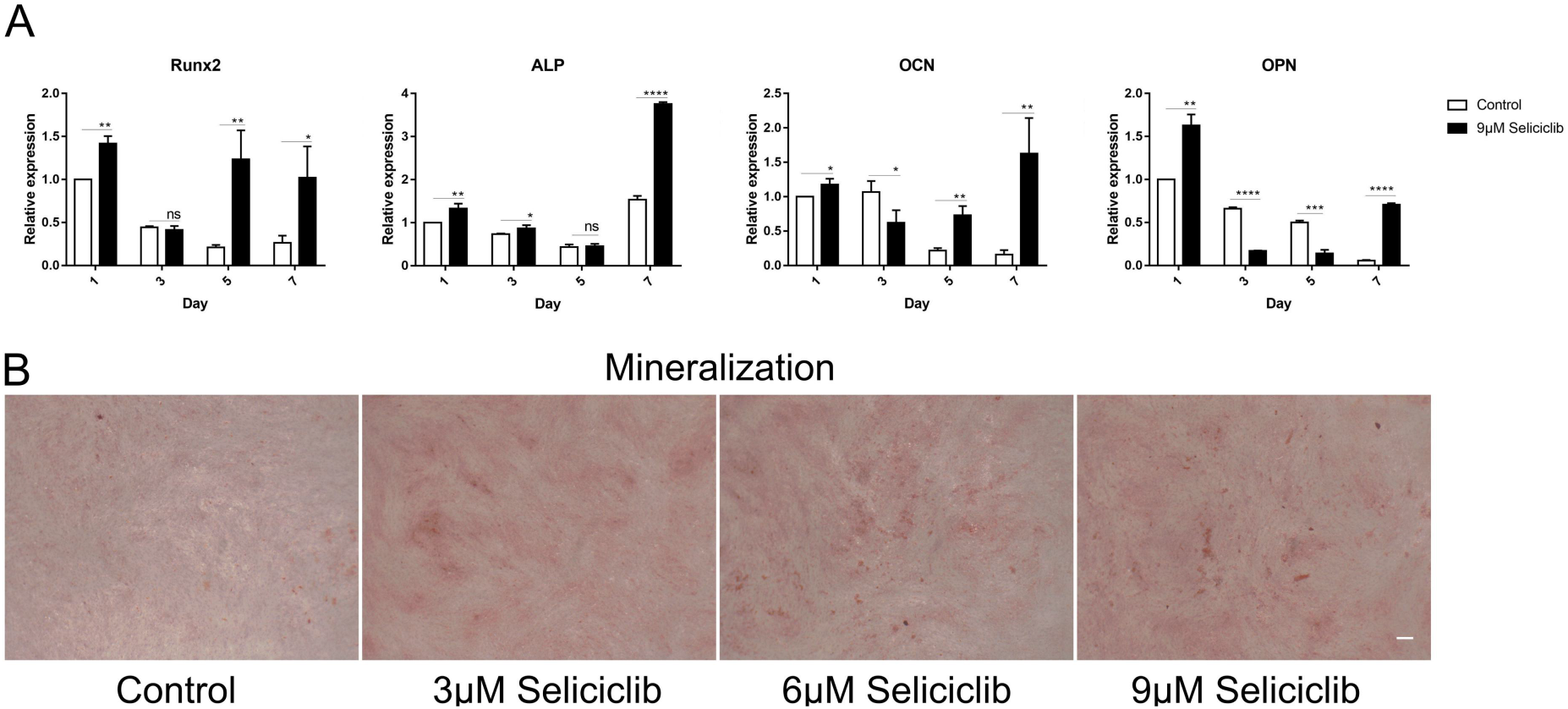
The expression of 4 osteoblastic differentiation marker genes of MSCs of rats and mineralization of MC-3T3E1 after Seliciclib treated. (A) The expression of Runx2, ALP, OCN and OPN (Left to right) between Seliciclib (9 μM) treated group (Black) and control (White) from day 1 to 7. Fold changes are relative to control of day 1. (B) Seliciclib promotes the mineralization of MC-3T3E1 at different concentrations of 3 μM, 6 μM and 9 μM. Scal bar 200 μm.

### Effects of Seliciclib on F-actin formation, nuclear localization of YAP and β-catenin, and RMRP RNA expression

Fluorescence microscopy revealed that, compared with the control group, the F-actin fibers formed in the Seliciclib-treated group appeared thicker, longer and more well-formed(Fig. 2. A). Moreover, the F-actin fibers in the Seliciclib-treated group exhibited more clear, whereas F-actin fibers were vague in the control group. Confocal microscopy showed that, in the Seliciclib-treated group, β-catenin was dominantly distributed in the nucleus (Fig. 2B). In comparison, in the control group, it appeared to be homogeneously distributed in the cell. The same tendencies were observed for YAP in the treated and control groups (Fig. 2C).RT-PCR found that, compared with the control, the expression of RMRP RNA in the Seliciclib-treated group was significantly lower on day 1 but was significantly higher on day 3, 5 and 7 (Fig. 2B). Furthermore, the intragroup difference in the expression of RMRP RNA increased between days 1–7.

**Figure 2.**
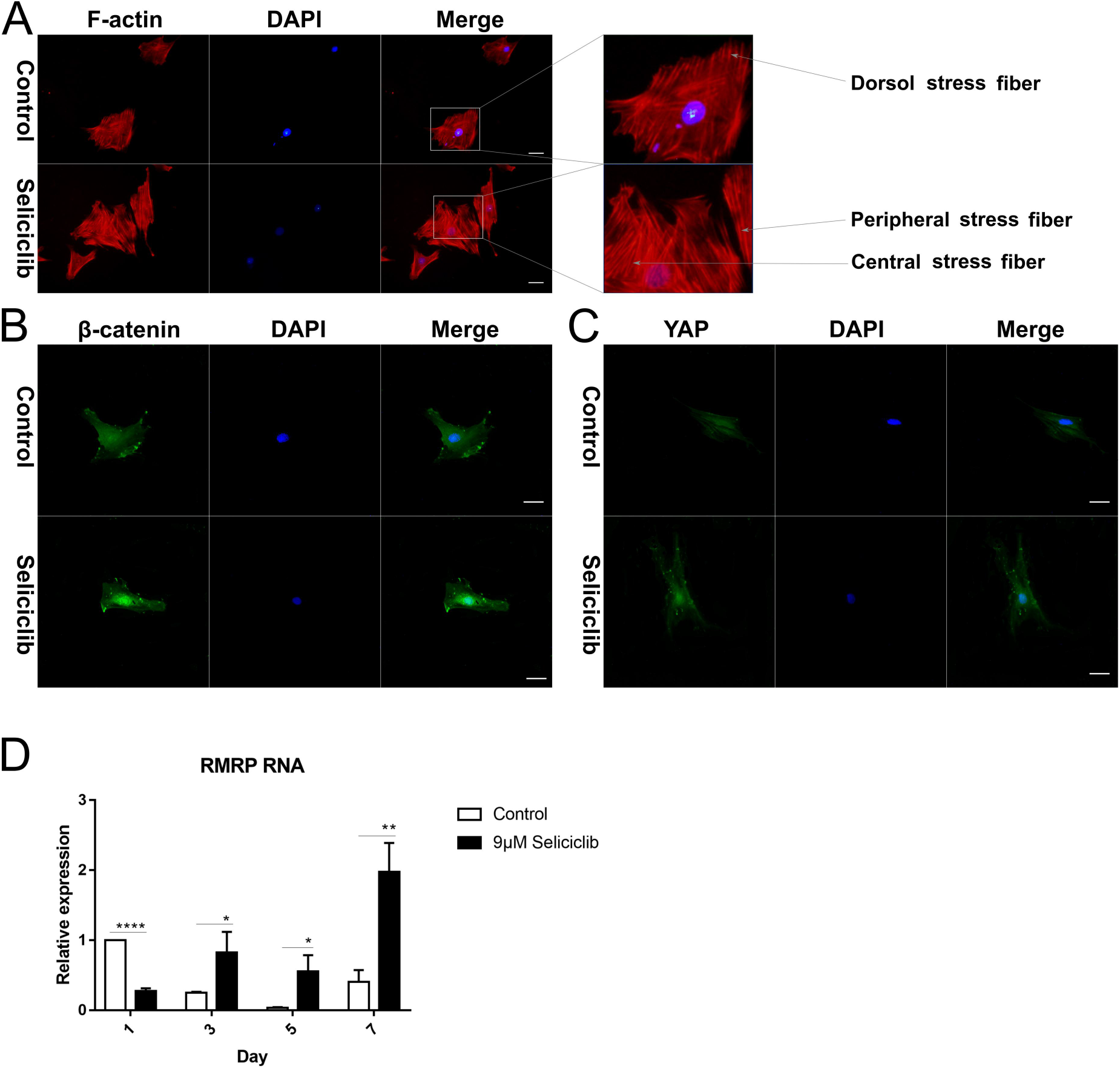
The F-actin development, YAP and β-catenin nuclear localization and the expression of RMRP RNA after Seliciclib-treated and control group. (A) The F-actin staining shows the development of Seliciclib (9 μM) -treated group (Above) and control (Below). Scal bar 50 μm. (B and C) Immunofluorescent staining of YAP and β-catenin after Seliciclib (9 μM) treated (Below) and control group (Above). Scal bar 25 μm. (D) RMRP RNA expression of Seliciclib-treated group (Black) and control (white) from day 1 to day 7. Fold changes are relative to control of day 1.

### Effects of F-actin inhibition on nuclear localization of YAP and β-catenin and expression of RMRP RNA

In the control group (treated with Seliciclib alone), many well-formed F-actin fibers were observed (Fig. 3A). In comparison, in the group simultaneously treated with Blebbistatin and Seliciclib, actin was distributed throughout the cell with the formation of a small number of fibers. Additionally, in the control group, YAP and β-catenin exhibited evident nuclear localization (Fig. 3B, C), whereas such trends were not found in the simultaneously treated group. Interestingly, the MSCs in the control group appeared spindle-like, whereas those in the simultaneously treated group were branched, resembling neurons.The expression levels of *RMRP RNA*, *Runx2*, and *ALP* in the simultaneously treated group were significantly lower than those in the control group (Fig. 3D), indicating that the down-regulation of CDK5 promoted *RMRP RNA* expression and osteoblastic differentiation at least via facilitating the development of stress fiber.

**Figure 3.**
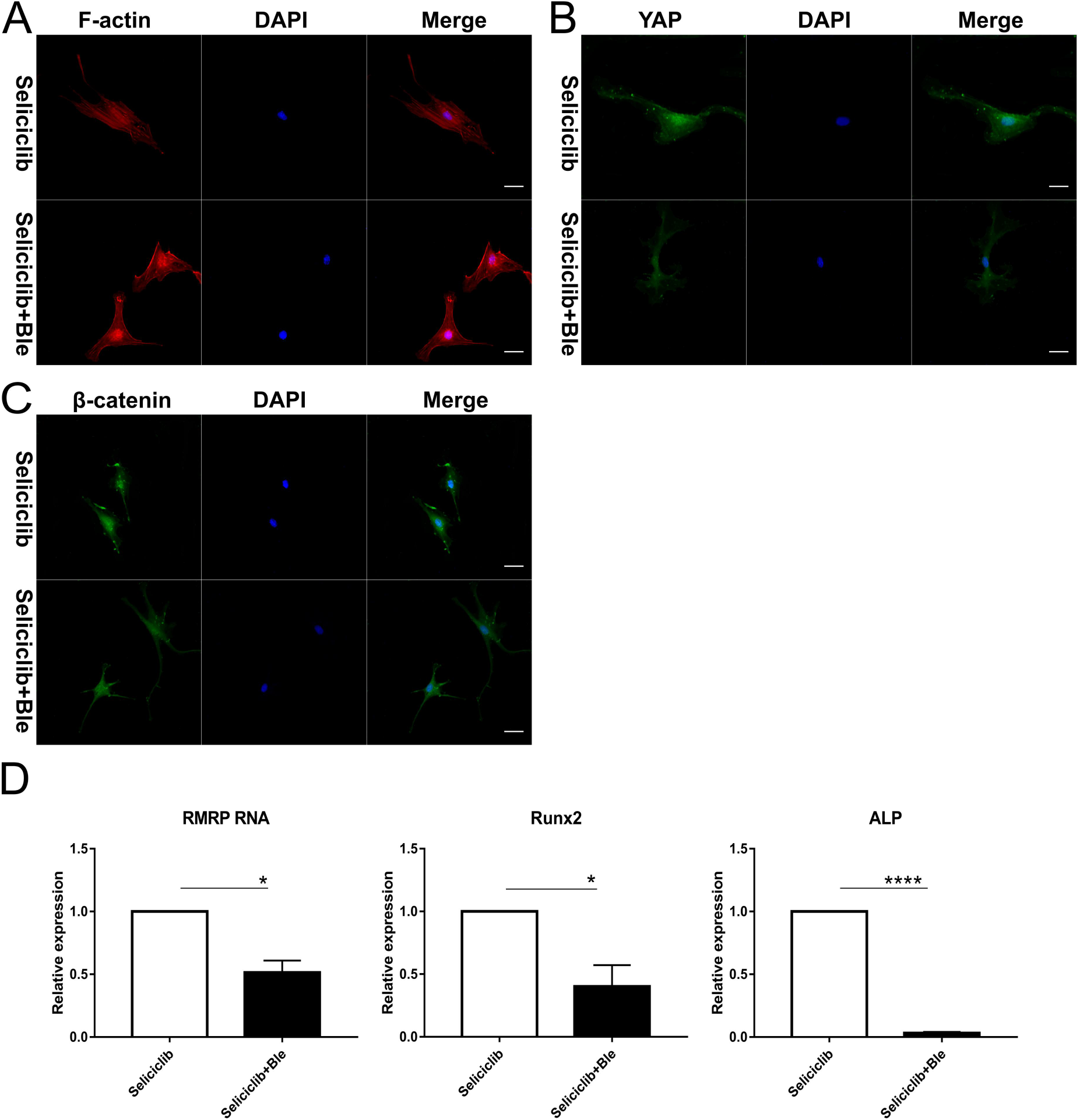
YAP and β-catenin nuclear localization and expression of *RMRP RNA*, *Runx2* and *ALP* after F-actin impressed. (A) F-actin development of Seliciclib (9 μM) -treated group (Above) and Seliciclib (9 μM) /Ble (5 μM) -treated group (Blow). Scal bar 25 μm. (B and C) Immunofluorescent staining of YAP and β-catenin shows their nuclear localiztion between Seliciclib (9 μM) -treated group (Above) and Seliciclib (9 μM) /Ble (5 μM) -treated group (Blow). Scal bar 25 μm. (D) Expression of *RMRP RNA*, *Runx2* and *ALP* of Seliciclib (9 μM) -treated group (white) and Seliciclib (9 μM) /Ble (5 μM) -treated group (Black). Fold changes are relative to Seliciclib-treated group.

### Effect of Seliciclib on migration of MG-63 cells

Crystal violet staining showed that, Seliciclib treatment reduced the migration of MG-63 cells in a concentration-dependent manner (Fig. 4A). Analysis of 5 randomly selected fields of view found that, compared with the control, the number of migratory cells decreased 25%, 45%, and 55% at 3 μM, 6 μM, and 9 μM, respectively (Fig. 4B). The difference between any two groups was statistically significant.

**Figure 4.**
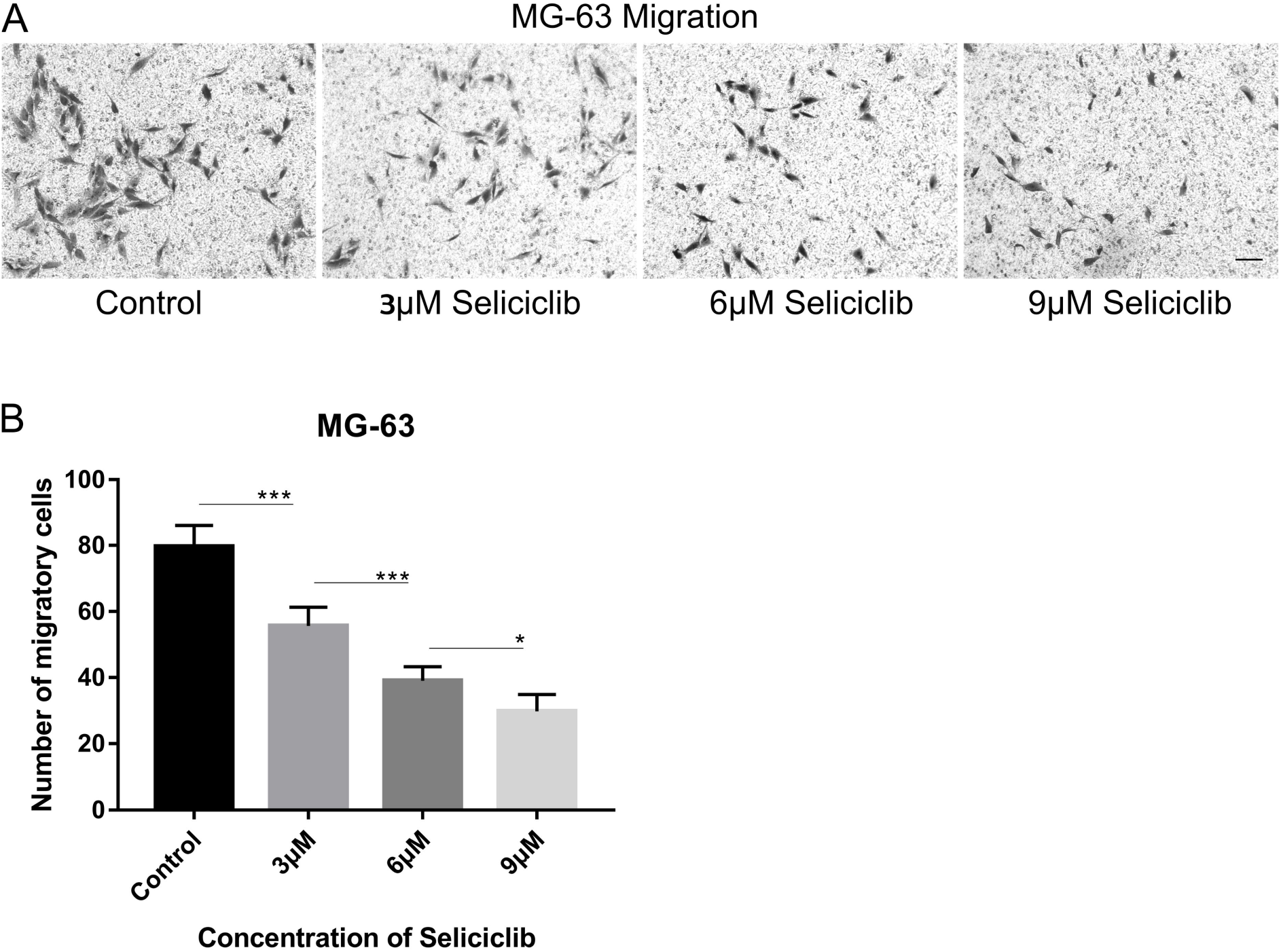
Migration of MG-63 cells after Seliciclib treated. (A) Crystal violet staining shows the migration of MG-63 of Seliciclib (3 μM, 6 μM and 9 μM) -treated group (Three on the right) and control (Left). Scal bar 100 μm. (B) Bar graph shows average migratory cells from Five randomly selected areas of different concentrations of Seliciclib (0, 3 μM, 6 μM and 9 μM from left to right). Date represents mean ± SEM.

## 4. Discussion

This study shows that CDK5 has a negative effect on F-actin formation of MSCs and CDK5 inhibition can promote their osteoblastic differentiation. Moreover, CDK5 indirectly modulates the β-catenin nuclear localization by suppressing the development of F-actin. Finally, we observed that CDK5 inhibition can suppress the migration of MG-63 cells. Together, these results suggest new potential functions of Seliciclib to promote the osteoblastic differentiation of MSCs and suppress the migration of osteosarcoma cells.

Inhibition of CDK5 may enhance F-actin development via two pathways. First, inhibited CDK5 level reduces the phosphorylation of Pak1 on T212, resulting in Rac1 activation [2, 3]. Additionally, CDK5 inhibition may reduce the phosphorylation of DLC1 on S120, S205, S422, and S509 sites, leading to its inactivation [4]. Because DLC1 is a GAP that can down-regulate the activation of RhoA [20, 21], CDK5 inhibition activates RhoA. Combined with myosin II, F-actin form stress fibers, which are categorized to dorsal, ventral, and peripheral fiber [22]. In the early stage of cell adhesion, activated Rac1 promotes the formation of the leading edge through generating a flat lamella, in which dorsal stress fiber exists. Dorsal stress fibers often lack myosin II, indicating that they are not contractile [23, 24]. Subsequently, RhoA is activated to facilitate the development (i.e., formation-thickening) of stress fibers and generation of a stress via RhoA-ROCK-RLCs signaling [25, 26].

In the present study, the CDK5 inhibition created thicker and longer stress fibers (vs. control) (Fig.2A). Moreover, most of the stress fibers formed in the Seliciclib-treated group were ventral and peripheral fibers. These suggest that, in the Seliciclib-treated group, stress fibers may be formed primarily via the second pathway. However, as antibodies to phosphorylated DLC1 on site S120, S205, S422, or S509 are not commercially available so far, this suggestion remains to be confirmed.

RMRP RNA, a long non-coding RNA, is a component of small nucleolar ribonucleoprotein (snoRNP) particle named RNase MRP. Mutations of RMRP RNA at different sites are sources of skeletal dysplasias such as cartilage-hair hypoplasia [27, 28]. Reduced RMRP RNA can down-regulate *Runx2,* and *ALP*,, and impair mineralization of bone-related cells, indicating RMRP RNA to be a positive regulator of osteoblastic differentiation [29]. However, molecular mechanisms underlying how RMRP RNA promotes osteoblastic differentiation are unknown. Beta-catenin and YAP promote the expression of RMRP RNA, both separately and synergistically [30]. CDK5/p35 complex can combine with β-catenin and then reduce N-cadherin-mediated and calcium-dependent cell-cell adhesion [8]. Additionally, CDK5/p35 complex combines with, and thus inactivates, β-catenin, a transcription factor able to stimulate osteoblastic differentiation [9]. In the current study, CDK5 inhibition by Seliciclib treatment increased nuclear localization of YAP and β-catenin and upregulated *RMRP RNA* expression (vs. control). Additionally, when CDK5 and F-actin were both inhibited, the nuclear localization of β-catenin was reduced (Fig. 3C), indicating that β-catenin nuclear localization to be F-actin-dependent. YAP plays an important role in maintaining the nuclear localization of β-catenin during osteoblastic differentiation [11]. Thus, after CDK5 inhibition, β-catenin is released. Meanwhile, nuclear localization of β-catenin might be enhanced by the increased nuclear localization of YAP. And these indicate CDK5 to be a upstream and negative regulator of RMRP RNA (Fig. 5).

**Figure 5.**
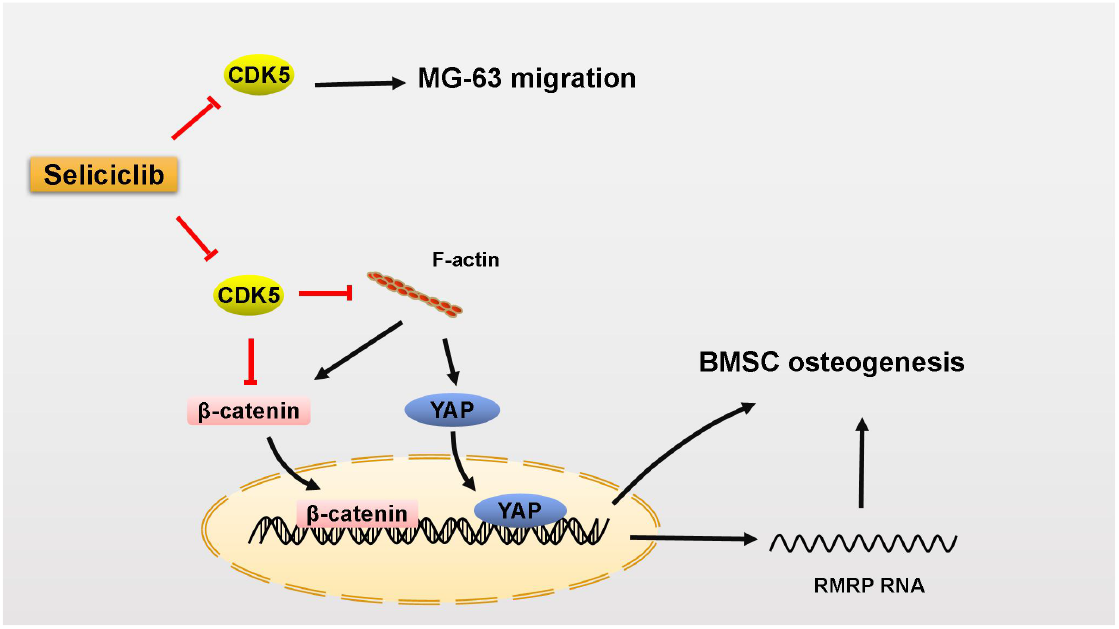
The model of Seliciclib promoting osteoblastic differentiation of MSCs and inhibiting the migration of MG-63. CDK5 inhibition releases β-catenin and promotes F-actin formation. Then well-formed F-actin fibers promote the nuclear localization of β-catenin and YAP, subsequently promoting the osteoblastic differentiation of MSCs partially by upregulating the expression of *RMRP RNA.* Moreover, CDK5 inhibition suppresess the migration of MG-63.

CDK5 inhibitor Seliciclib can promote osteoblastic differentiation. The inhibition of MG-63 cell migration elevated when Seliciclib concentration was increased from 3 to 9 μM. This suggests that, Seliciclib may have potential functions in bone defect repair, especially when suppression of osteosarcoma metastasis is required simultaneously. The amount of Seliciclib used to treat cancer in clinical trials is far greater than the amount used to promote osteoblastic differentiation, and the target of Seliciclib used to treat cancer is mainly CDK1/2 or ERK1/2 [31]. The IC50 of Seliciclib for CDK5/P35, CDC2/cyclinB, CDk2/cyclinA, ERK1 and ERk2 in vitro is 0.16, 0.65, 0.7, 34 and 14 μM respectively [32].

Although these results reveal that, the down-regulation CDK5 promotes the osteoblastic differentiation of MSCs partially by enhancing the development of F-actin, the mechanisms responsible for CDK5 down-regulation are unknown and need to be identified. CDK5, RhoA, YAP, β-catenin and RMRP RNA play an important roles in the cancer cell migration, invasion, proliferation and morphological change of tumor cells [16, 21, 33–35]. Results of this study suggest a link between CDK5, RhoA, YAP, β-catenin, and RMRP RNA and new insights for understanding the molecular mechanisms of some carcinogenesis.

## Acknowledgement

The research was supported by National Natural Science Foundation of China (32071343), Fundamental Research Funds for the Central Universities (2682020ZT80) and Applied Basic Research Project of Sichuan Province (21YYJC3323).

## Author Contributions

H.F. designed the experiments; T.L.G. and H.F. generated the idea and T.L.G. supervised the project; H.F. and K.D. wrote the manusctip; H.F. performed the most of the experiments with the supervision of T.L.G. H.Y.Z., R.F.W. and Y.L.Y. assisted with experiments.

